# A generative model of the hippocampal formation trained with theta driven local learning rules

**DOI:** 10.1101/2023.12.12.571268

**Authors:** Tom M George, Caswell Barry, Kimberly Stachenfeld, Claudia Clopath, Tomoki Fukai

## Abstract

Advances in generative models have recently revolutionised machine learning. Meanwhile, in neuroscience, generative models have long been thought fundamental to animal intelligence. Understanding the biological mechanisms that support these processes promises to shed light on the relationship between biological and artificial intelligence. In animals, the hippocampal formation is thought to learn and use a generative model to support its role in spatial and non-spatial memory. Here we introduce a biologically plausible model of the hippocampal formation tantamount to a Helmholtz machine that we apply to a temporal stream of inputs. A novel component of our model is that fast theta-band oscillations (5-10 Hz) gate the direction of information flow throughout the network, training it akin to a high-frequency wake-sleep algorithm. Our model accurately infers the latent state of high-dimensional sensory environments and generates realistic sensory predictions. Furthermore, it can learn to path integrate by developing a ring attractor connectivity structure matching previous theoretical proposals and flexibly transfer this structure between environments. Whereas many models trade-off biological plausibility with generality, our model captures a variety of hippocampal cognitive functions under one biologically plausible local learning rule.

## 1 Introduction

Generative models seek to create new data samples which are similar to those from the training set. To do so they must learn the probability distribution of the training data, comprising a rich, generalisable and accurate model of the world. Many of the recent advances in AI have involved types of generative models: VAEs [1], GANs [2], diffusion models [3] and autoregressive models [4] have seeded improvements in AI capabilities ranging from data compression [5] to image generation [6] and natural language [7]. In neuroscience, the animal brain has long been known to exploit generative models [8, 9]. The ability to generate representative sensory data samples can be used directly, for example during offline planning or memory recall. It can also be used indirectly to aid training of inference networks with the goal of processing rich, noisy and high dimensional streams of incoming sensory stimuli, as discussed in the predictive coding literature [10]. In a sentence: “What I cannot create [generate], I do not understand [inference]” (R. Feynman).

The hippocampal-entorhinal system (aka. hippocampal formation) – a brain structure implicated in spatial [11] and non-spatial [12] memory – provides a pertinent example. Its primary role seems to be inference [13]: mapping sensory inputs into a robust and decodable representation of state (grid cells [14], place cells [11] etc. [15]). A generative model is thought to have a dual role in learning: supporting offline tasks such as route planning [16] and memory consolidation [17], and online during behaviour with path integration [18]. Path integration enables the hippocampal network to maintain an up-to-date and accurate estimate of its position in the absence of reliable sensory data by integrating self-motion cues. A recent flurry of computational [19, 20, 21] and theoretical [22, 21] work has highlighted the importance of path integration as a key objective explaining hippocampal function and representations.

Existing computational generative models of the hippocampal formation [23, 24] account for many of its cognitive functions and internal representations but require non-trivial learning rules and message passing protocols which don’t connect with known aspects of biology. Computational models of path integration [25, 26, 27] have mostly focussed on continuous attractor networks which, although experimentally supported [28], alone lack the complexity or expressivity required of a fully general model of the hippocampal memory system.

The primary contribution of this paper is to introduce a biologically plausible model of sequence learning in the hippocampus which unifies its capacities as a generative model of sensory stimuli and path integration under one schema. To do this we propose modeling the hippocampal formation as a Helmholtz machine [29] which learns to predict sensory stimuli given the current hidden state and action (e.g. velocity). We propose a deep connection between the hippocampal theta oscillation [30] and the unsupervised wake-sleep algorithm [31] for training Helmholtz machines. Though this class of generative models isn’t widely used, and lacks the scalability of the lastest transformer-based sequence learners, it excels in this context since is has many natural points of contact with biology (both in terms of architecture and neural dynamics) yet still maintains the expressiveness afforded to models of the brain by deep neural networks.

In this paper we:

- introduce a new model of the hippocampal formation which learns the latent structure of an incoming stream of sensory stimuli analogous to a Helmholtz machine.
- describe a biologically plausible learning regime: Theta-oscillations gate information flow through multi-compartmental neurons which rapidly switches the system between “wake” and “sleep” phases. All plasticity is local.
- train our model on stimuli from a biologically relevant spatial exploration task and show it learns to path integrate by developing a ring attractor connectivity structure (comparable to theoretical predictions and empirical results in deep recurrent neural networks trained with gradient descent). Learning generalises: when the agent moves to a new environment, path integration capabilities recover without needing to relearn the path integration weights.

Our model of the hippocampal formation simultaneously (i) accounts for its role as a generative model of sensory stimuli, (ii) can learn to path integrate and (iii) can transfer structural knowledge between environments. The model, though here applied to the hippocampus, can be viewed as a step towards a general solution for how biological neural networks in many brain regions (for example visual cortex [10]) can learn generative models of the world.^1^

### 1.1 Related work

A recent generative model of the hippocampus, the Tolman-Eichenbaum Machine [23], proposed that the hippocampal formation be thought of as a hierarchical network performing latent state inference. Medial entorhinal cortex (MEC) sits atop the hierarchy and learns an abstract representation of space which is mapped to the hippocampus (HPC) where it is bound onto incoming sensory stimuli. Once trained the system can act in a generative fashion by updating the hidden representation with idiothetic action signals and then predicting the upcoming sensory experience. The drawback of this model, and others which share a similar philosophical approach [32, 24], is that it requires training via backpropagation through time (or equivalent end-to-end optimisation schemes, as in [24]) without clear biological correlates. Related hierarchical network architectures have also been studied in the context of reinforcement learning [33] and hippocampal associative memory [34].

Historically, hippocampal models of path integration have focused on continuous attractor networks (CANs) [25, 26, 27, 21] in entorhinal cortex. A bump of activity representing location is pushed around the CAN by speed and/or head-direction selective inputs, thus integrating self-motion. CANs have received substantial experimental support [28] but few studies adequately account for *how* this structure is learned by the brain in the first place. One exception exists outside the hippocampal literature: Vafidis et al. [35] built a model of path integration in the fly head-direction system which uses local learning rules. Here we go further by embedding our path integrator inside a hierarchical generative model. Doing so additionally relaxes the assumption (made by Vafidis et al. [35] and others [36]) that sensory inputs into the path integrator are predefined and fixed. Instead, by allowing all incoming and outgoing synapses to be learned from random initalisations, we achieve a more generalisable model capable of transferring structure between environments (see section 3.3).

Hippocampal theta oscillations have been linked to predictive sequence learning before [37, 38, 39, 40] where research has focused on the compressive effects of theta *sequences* and how these interplay with short timescale synaptic plasticity. Instead of compression, here we hypothesize the role of theta is to control the direction information flows through the hierarchical network.

Finally, a recent theoretical work by Bredenberg et al. [41] derived, starting from principles of Bayesian variational inference, a biologically plausible learning algorithm for approximate Bayesian inference of a hierarchical network model built from multi-compartmental neurons and trained with local learning rules using wake-sleep cycles. Here we build a similar network to theirs (i) extending it to a spatial exploration task and mapping the hidden layers onto those in the hippocampal formation, (ii) simplifying the learning rules and relaxing a discrete-time assumption – instead, opting for a temporally continuous formulation more applicable to biological tasks such as navigation – and (iii) adapting the hidden layer to allow idiothetic action signals to guide updates (aka. path integration). Their work provides a theoretical foundation for our own, helping to explaining *why* learning converges on accurate generative models.

## 2 A biologically plausible generative model trained with rapidly switching wake-sleep cycles and local learning rules

In sections 2 and 3 we give concise, intuitive descriptions of the model and experiments; expanded details can be found in the supplementary material.

### 2.1 Basic model summary

We consider learning in an environment defined by a latent state, *z*(*t*), which updates according to stochastic dynamics initially unknown to the network,

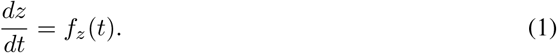

These dynamics depends on the task; first we consider *z*(*t*) to be a set of mutually independent random variables and later we consider the more realistic task of an agent moving on a 1D track.

The network recieves sensory input which is a function of the latent state into a sensory layer, **p**(*t*), and communicates this to a hidden layer (aka “internal state”), **g**(*t*). The network contains both an *inference* (aka. *recognition*) model which infers the hidden state from the sensory input (green arrows, Fig. 1a) and a *generative* model which updates the hidden state with recurrent synapses and maps this back to the sensory layer (blue arrows). As we will soon identify these processes with Basal and Apical dendritic compartments of pyramidal neurons we label activations sampled from the inference model with the subscript *B* and those from the generative model with the subscript *A*. ^2^ In summary

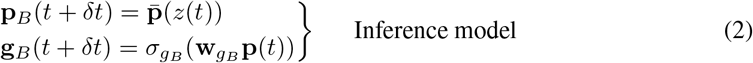

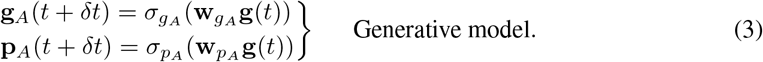

**w**_*g B*_, **w**_*p A*_, **w**_*g B*_ are matrices of randomly initialised and plastic synaptic weights. 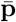 maps the environmental latent into a vector of neural inputs. *σ*’s denote activation functions applied to the dendritic pre-activations – either the identity (*σ*(*x*) = *x*) or rectified tanh functions (*σ*(*x*) = max(0, *tanh*(*x*))). A small amount of noise is added to the dendritic activations to simulate realistic biological learning.

**Figure 1:**
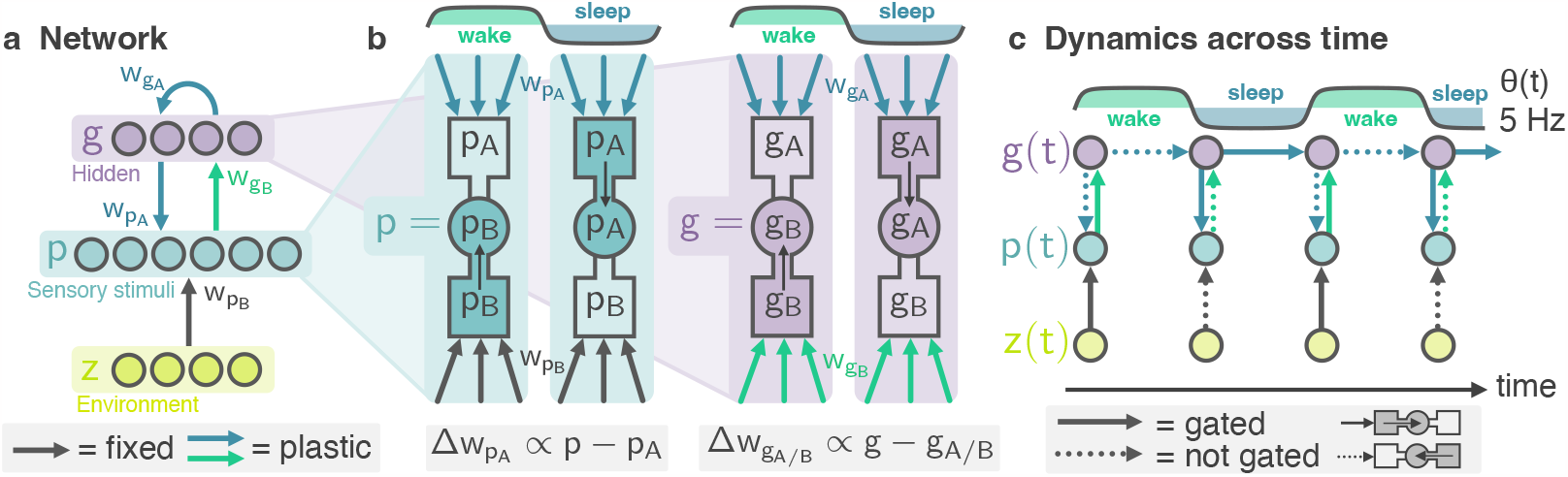
A biologically plausible generative model is trained with theta frequency wake-sleep cycles and a local learning rule. **a** Network schematic: high-D stimuli from an underlying environmental latent, *z*, arrive at the basal dendrites of the sensory layer, *p*, and map to the hidden layer, *g* (this is the inference model, weights in green). Simultaneously, top-down predictions from the hidden layer *g* arrive at the apical dendrites of *p* (this is the generative model, weights in blue). **b** Neurons in layers *p* and *g* have three compartments. A fast oscillation, *θ*(*t*), gates which dendritic compartment – basal (*p*_*B*_, *g*_*B*_) or apical (*p*_*A*_, *g*_*A*_) – drives the soma. A local learning rule adjusts input weights to minimise the prediction error between dendritic compartments and the soma. **c** This equates to rapidly switching “wake” and “sleep” cycles which train the generative and inference models. Panel c displays just two updates per theta-cycle, in reality there are many (*δt << T*_*θ*_).

We believe that the widely adopted convention of modelling neurons as single-compartment perceptrons is limiting. By considering, in a minimal extension, the distributed dendritic structure of real neurons we can tap into significant potential for explaining hippocampal learning. Theoretical [42, 43, 44, 45] and experimental [46, 47, 48] research into credit assignment in biological neurons has identified different roles for basal and apical dendrites: basal dendrites are thought to receive bottom-up drive from sensory inputs whereas apical dendrites receive top-down drive from higher layers in the sensory hierarchy [49]. Following this line of research — and matching an equivalent theoretical model of latent state inference described by [41] — we identify the inference process with synaptic inputs into a basal dendritic compartment of pyramidal neurons and the generative process with synaptic inputs into an apical dendritic compartment. In summary, each **p** and **g** neuron in our model has three compartments: a somatic compartment, a basal dendritic compartment and an apical dendritic compartment (Fig. 1b). Only the somatic activation is used for communication between layers (right hand side of Eqns. (2) and (3)) while dendritic compartment activations are variables affecting internal neuronal dynamics and learning as described below (Eqns. (4) and (6)).

### 2.2 Theta oscillations gate the direction of information flow through the network

The dynamics of the somatic activations **p**(*t*) and **g**(*t*) are as follows: the voltage in each soma is either equal to the voltage in the basal compartment *or* the voltage in the apical compartment depending on the phase of an underlying theta oscillation. This is achieved by a simple theta-gating mechanism (Fig. 1b):

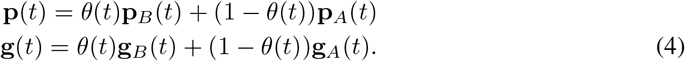

where *θ*(*t*) is a 5 Hz global theta oscillation variable defined by the square wave function:

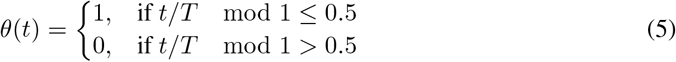

for *T* = 1*/f*_*θ*_ and *f*_*θ*_ = 5 Hz, matching the hippocampal theta frequency (5-10 Hz) [50]. According to this model theta-band oscillations in the hippocampal local field potential gate which dendritic compartment drives the soma. Experimental [47, 51, 52] and modelling work [53] gives provisional support for this assumption.

These local theta-dynamics have global consequences: the early phase (*θ*(*t*) = 1) of each theta cycle can be thought of as a “wake” phase where information flows upwards through the network from the environment to the hidden layer, sampling the inference model. The latter phase (*θ*(*t*) = 0) of each theta cycle is a “sleep” phase where information flows down from the hidden layer to the sensory units, sampling the generative model. These dynamics are displayed in Fig. 1.

### 2.3 Hebbian-style learning rules train synapses to minimise local prediction errors

In contrast to comparable models which are optimised end-to-end using backpropagation through time our model learns synaptic weights according to a local plasticity rule which is a simplified variant of a rule proposed by Urbanczik and Senn [43]. Incoming synaptic projections are continually adjusted in order to minimize the discrepancy between the somatic activation and the dendritic activation. The full learning rules are described in the supplement but simplified versions are given here:

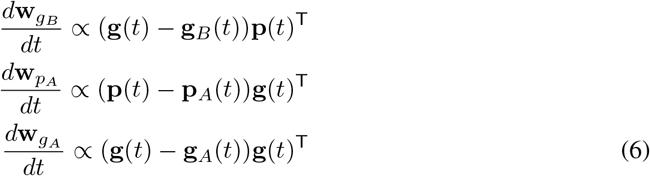

Notably this learning rule is equivalent for *all* plastic synapses in the model: **p** to **g, g** to **p** and the recurrent **g** to **g** synapses (see Fig. 1b). If a local prediction error is detected, for example the somatic activation is larger than the dendritic activation, then the synaptic strength of inputs into that dendritic compartment which are positive/negative are strengthed/weakened to reduce the error. This model can equivalently be viewed as a type of Hebbian learning – weight change is proportional to the correlation of pre- and post-synaptic activity (the first term) – regularised (by the second term) to prevent unbounded growth.

During the wake phase the weights of the generative model (**w**_*p*_ and **w**_*g A*_ ) are trained and plasticity on the inference weights (**w**_*g B*_ ) falls to zero. This occurs naturally because **p** = **p**_*B*_ so there will be no basal prediction errors to correct. During sleep the reverse occurs; the weights of the inference model are trained and plasticity on the generative model falls to zero. Experimentally, apical activity is known to guide plasticity at basal synapses in CA1 [46]. This alternating, coordinated regime of sampling and learning (sample-inference-train-generative, then sample-generative-train-inference) is a hallmark of the wake-sleep algorithm. It fundamentally differs from the forward and backward sweeps of backpropagation since neurons remain provisionally active at all times so the process of learning minimally perturbs perception. Also, whereas backpropagation sends error signals down through the network to train synaptic weights, here only predictions are sent between layers and error signals are calculated locally at each dendrite.

As discussed in section 1, Bredenberg et al. [41] mathematically derive learning rules similar to these starting from a loss function closely related to the evidence lower bound (ELBO). As such our identification of early- and late-theta phases as “wake” and “sleep” cycles can be considered precise: from a Bayesian perspective our hippocampal model is minimising a modified ELBO loss (see supplement) thus learns to find approximately optimal inference and generative models accounting from the temporally varying stimulus stream it is presented.

### 2.4 Velocity inputs into the hidden layer

For path integration, the hidden state needs access to an idiothetic (internally generated) velocity signal. To satisfy this we endow the hidden layer, **g**, with conjunctive velocity inputs, henceforth “conjunctive cells”, as shown in Fig. 3a & b. Conjunctive cells are organised into two groups: **g**_*v*_*L* is responsible for leftward motion and **g**_*v R*_ for rightward motion. Each conjunctive cell receives input from the hidden units and either the leftward (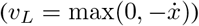 or rightward 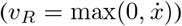 ) component of the velocity. For the results shown this connectivity is one-to-one [**w**_*gvL*_ ]_*ij*_ = [**w**_*gvR*_ ]_*ij*_ = *δ*_*ij*_ but random connectivity works too, see supplement. Finally, conjunctive cells send return connections back to the apical dendritic compartment of the hidden units via a randomly initialised plastic synaptic weight matrix. This inputs are what drive the hidden units to path integrate.

This model takes inspiration from so-called conjunctive grid cells [54] found in the medial entorhinal cortex (MEC). These cells, though to be an integral component of the mammilian path integration system[27], are jointly tuned to head direction and location much like the conjunctive cells in our model. An important and novel aspect of our model is that synaptic weights between or into the hidden units are *learned*. This deviates from other models for example that by Burak and Fiete [27] (where all connectivity is predefined and fixed) or Vafidis et al. [35] and Widloski and Fiete [36] (where sensory inputs to the hidden units are pre-defined and fixed). This is not only more realistic but affords the model flexibility to translate path integration abilities between environments without having to relearn them, a form of transfer learning which we demonstrate in section 3.3.

## 3 Results

### 3.1 Validation on an artifical latent learning task

We begin by testing the basic model (i.e. without conjunctive inputs, Fig. 1a) on an artificial task. *N*_*z*_ = 5 latents, *z*_*i*_(*t*), are independently sampled from a smooth, random process with an autocorrelation timescale of 1 second (Fig. 2a). The sensory layer, *N*_*p*_ = 50, then receives a high-dimensional random linear mixture of the latents into the basal compartments:

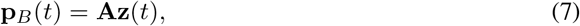

where **A** ∈ ℝ^50*×*5^ and 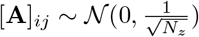. The hidden layer, **g**(*t*), is matched in size to the latent process, *N*_*g*_ = *N*_*z*_ = 5, and all dendritic activation functions are linear. We train the model for 30 minutes of simulated time and track prediction errors, the difference between the basal and apical activations in the sensory and hidden layers, which reliably decreased throughout training (Fig. 2b). We then perform two tests designed to confirm whether the model has learnt accurate inference and generative models.

**Figure 2:**
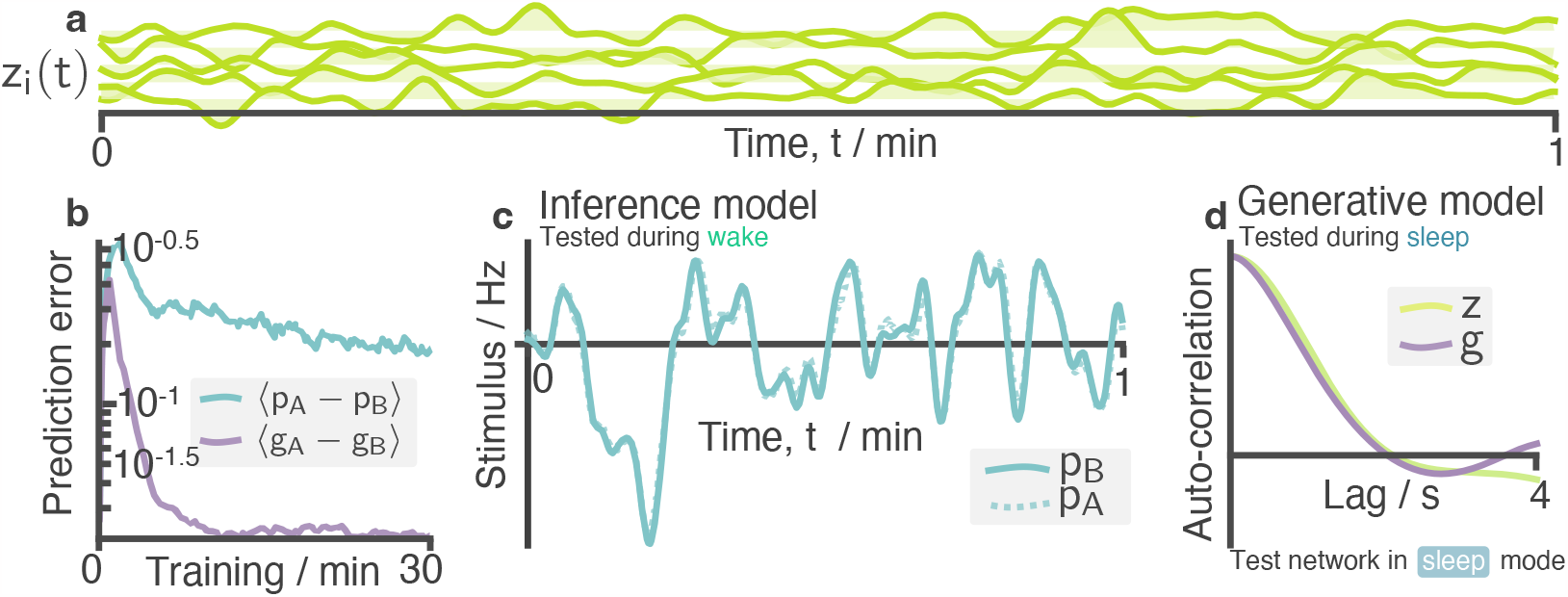
Learning in an environment of temporally varying latents. **a** In this artifical task the latent space comprises of *N*_*z*_ = 5 independent random variables with an autocorrelation decay timescale of 1 s. **b** Prediction errors (difference between apical and basal activations) in sensory and hidden layers reduce over training time. **c** Tested in wake mode (*θ* = 1) after training, the ground truth stimulus matches apical prediction for all stimulus dimensions (one shown) implying the network is efficiently “autoencoding” the sensory inputs into and back out of the compressed hidden layer. **d** Tested in sleep mode (*θ* = 0, no environmental inputs), generated data from the hidden units, *g*, have an autocorrelation curve which matches that of the true latents implying a statistically accurate generative model has been learned. More extensive samples from this model, before and after training, can be found in Fig. S1

**Figure 3:**
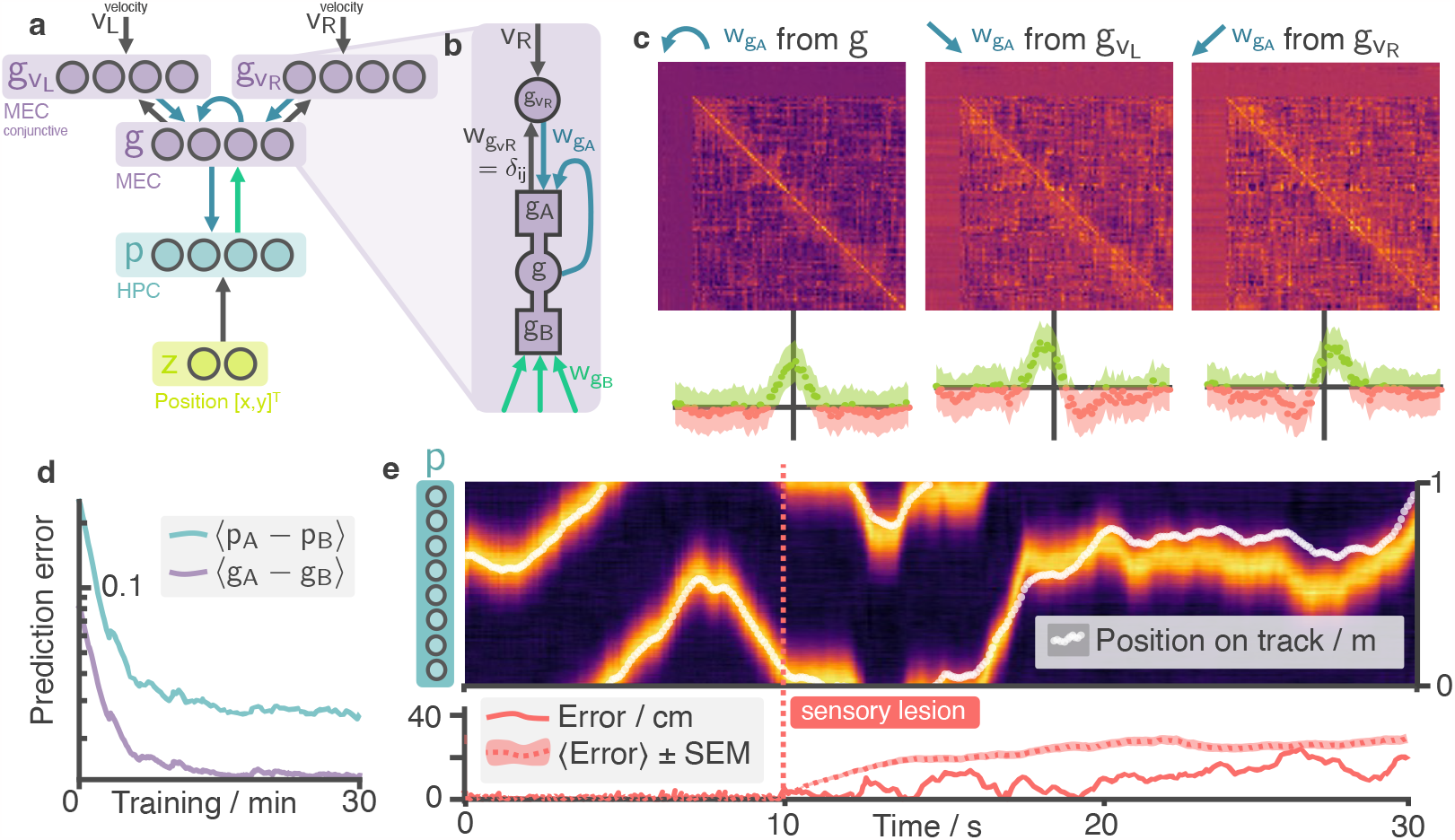
The hippocampal model learns to path integrate on a 1D track using a ring attractor. **a** Position selective (place cell) inputs drive basal dendrites of the sensory layer **p** (HPC). **b** Hidden units (MEC) are connected to two sets of “conjunctive cells” which each connect back to one of the hidden neurons (**g**) and either the leftward (for **g**_*v L*_ ) or rightward (for **g**_*v L*_ ) velocity of the agent allowing velocity information to enter the network. Synaptic strengths of the return connections from the conjunctive cells to the MEC hidden units, as well as those for the MEC recurrent connectivity (collective denoted **w**_*g A*_ ), are randomly initialised and plastic. **c** After training, reordering the hidden units by the position of peak activity reveals a ring attractor in the synaptic weight matrices. Centre-surround recurrent connectivity stabilises an activity bump which is then “pushed” around the attractor manifold by asymmetric connections from the conjunctive cells, integrating velocity. Bands of zero weights show MEC neurons which have become perpetually inactive (aka “died”). The bottom panel displays the matrix row-averages, utilizing the circular symmetry of the environment to align rows before averaging. **d** Learning plateaus after 15 mins of simulated time. **e** Path integration ability is demonstrated in a lesion study: after 10 seconds in the normal oscillatory mode the network is placed into sleep mode (aka generative mode), lesioning the position-dependent sensory inputs. Despite this HPC continues to accurately encode position, evidence that the MEC ring attractor is path integrating the velocity inputs and sending predictions back to HPC. Lower panel shows the accumulated decoding error as well as the mean±SEM over 50 trials.

First, we set the dynamics of the model to “wake” mode (*θ* = 1) and measure the basal and apical activations of one of the sensory neurons for 60 seconds. Close correspondence (Fig. 2c) confirms that the network accurately “autoencodes” the high-dimensional sensory inputs through the compressed hidden layer. Since all activation functions are linear this implies that **w**_*g B*_ and **w**_*p A*_ are pseudoinverses. Next, we place the network in “sleep” mode (*θ* = 0) and allow the generative model to run freely. The autocorrelation of the generated hidden states (**g**(*t*|*θ* = 0), displayed fully in the supplement) match that of the true environmental latents (**z**(*t*)), Fig. 2d, implying the generative model has statistics closely matching those of the true underlying generative process.

### 3.2 Learnable path integration with a hidden ring attractor

Next we turn our attention to the hippocampal formation’s role in spatial navigation, and our central result. The environment consists of an agent randomly moving around a 1 m 1D circular track (motion and cell data is generated using the RatInABox package [55]). The basal compartment of each HPC neuron is spatially tuned to a single different Gaussian input however non-Gaussian randomly spatially tuned inputs work as well (see supplement Fig. S2b):

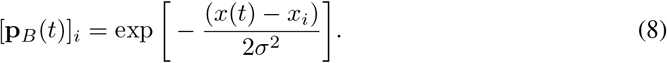

*x*(*t*) is the position of the agent and 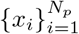 are the centres of the Gaussian inputs (*σ* = 6 cm), intended to simulate hippocampal place fields, evenly spaced at 1 cm intervals along the track. MEC (i.e. the hidden layer, **g**(*t*)) is matched in size *N*_*g*_ = *N*_*p*_ = 100 with rectified tanh activation functions on both dendritic compartments (*σ*_*g B*_ (*x*) = *σ*_*g A*_ (*x*) = max(0, tanh(*x*))) and HPC (the sensory layer **p**(*t*)) is linear (*σ*_*p*_*A* (*x*) = *x*). Two populations of conjunctive cells (Fig. 3a & b) feed into the apical compartments of the MEC recurrent units. Random initialisation of **w**_*g B*_ means that MEC neurons start off with random non-Gaussian spatial tunings. **w**_*g*_*A* and **w**_*pA*_ are also randomly initialised.

The network is trained for 30 minutes with learning plateauing after 15 (Fig. 3d). A lesion study, designed to test path integration, is then performed as follows: First, the network is run for 10 seconds normally (i.e. with theta-oscillating periods of wake and sleep). Since the simulated HPC neurons receive place-tuned inputs uniformly ordered along the track (i.e. *x*_*j*_ *> x*_*i*_∀*i, j > i*) an activity heatmap of HPC reveals a bump of activity accurately tracking agent’s position (Fig. 3e, left). The network is then placed into a sleep phase (*θ* = 0) for 20 seconds. This amounts to a full sensory lesion since top-down MEC inputs, not bottom-up place-tuned sensory inputs, drive HPC. Despite the full sensory lesion, hippocampal activity remains approximately unperturbed and the activity bump continues to accurately track position, slowly accumulating errors (Fig. 3e right). Since our HPC layer has no recurrent connectivity it cannot support this post-lesion activity on its own. Instead feed-forward drive from an MEC ring attractor, which we turn our attention to now, is responsible for maintaining the HPC code.

To find the ring attractor we must first reorder the MEC cells. We do this according to the position of the peak of their receptive fields (defined in the supplement). After reordering, the recurrent connectivity matrix can be seen to have acquired a centre-surround connectivity profile. Nearby MEC cells were, on average, strongly and positively recurrently connected to one another. Those far apart weakly inhibit one another (Fig. 3c, left; band of strong positive weights along diagonal flanked by weak negative weights). This profile matches that of a quasi-continuous ring attractor: local excitatory and long-range inhibitory connections stabilise a bump of activity on the attractor manifold in the absence of sensory input [56]. Weights from the conjunctive cells acquired asymmetric connectivity (Fig. 3c, middle & right) skewed towards the velocity direction for which they are selective. These asymmetric connections enable conjunctive cells to “push” the activity bump around the manifold, integrating velocity (see supplement for a visualisation of the MEC bump attractor). Theoretical work on ring attractors has demonstrated that for accurate path integration the asymmetric weights must be proportional to the derivative of the symmetric weights [56], approximately observed here. A noteworthy observation is that some MEC neurons become perpetually inactive; this is a consequence of the fact that *both* top-down and bottom-up synapses into the hidden layer are plastic and can fall to zero (Fig. 3c bands of zero-weights) satisfying a trivial *g*_*A*_ = *g*_*B*_ = 0 solution for minimising the prediction error. Despite this, not all MEC neurons die and the surviving subset are sufficient for path integration. In supplementary section 5.4.2 we discuss additional results showing when the network learns robust path integrate under a variety of plasticity, initialisation and noise manipulations.

Crucially, what sets this model apart from others [19, 20, 21, 22] is that the network is not optimized using a conventional path-integration objective and backpropagation. Instead, it has been demonstrated how path integration can naturally arise in a biologically constrained network subject to a much simpler (yet more broadly applicable) local objective, in cases where idiothetic velocity signals are available to the hidden layers.

### 3.3 Remapping: transfer of structural knowledge between environments

Finally, we demonstrate how our trained network can transfer structural knowledge – which here means the ring attractor and thereby path integration – between environments. We start by training the network as in section 3.2; the only diffence is that for simplicity we choose to fix **w**_*g B*_ = *δ*_*ij*_ giving rise to MEC representations which, like HPC, are unimodal (this constraint can be relaxed and, in the more general case, MEC units typically have multiple receptive fields, Fig S4d, reminiscent of grid cells). We then simulate a hippocampal “remapping” event by shuffling the sensory inputs to the HPC layer (Fig. 4a & b, top panel) and retraining the network for a further 30 minutes but this time holding weights in the hidden layer, **w**_*gA*_ . Only the HPC ↔ MEC synapses (**w**_*gB*_ *&* **w**_*pA*_ ) remain plastic during retraining. Biologically this may be accounted for by the observation that cortical plasticity is substantially slower than hippocampal plasticity [57].

**Figure 4:**
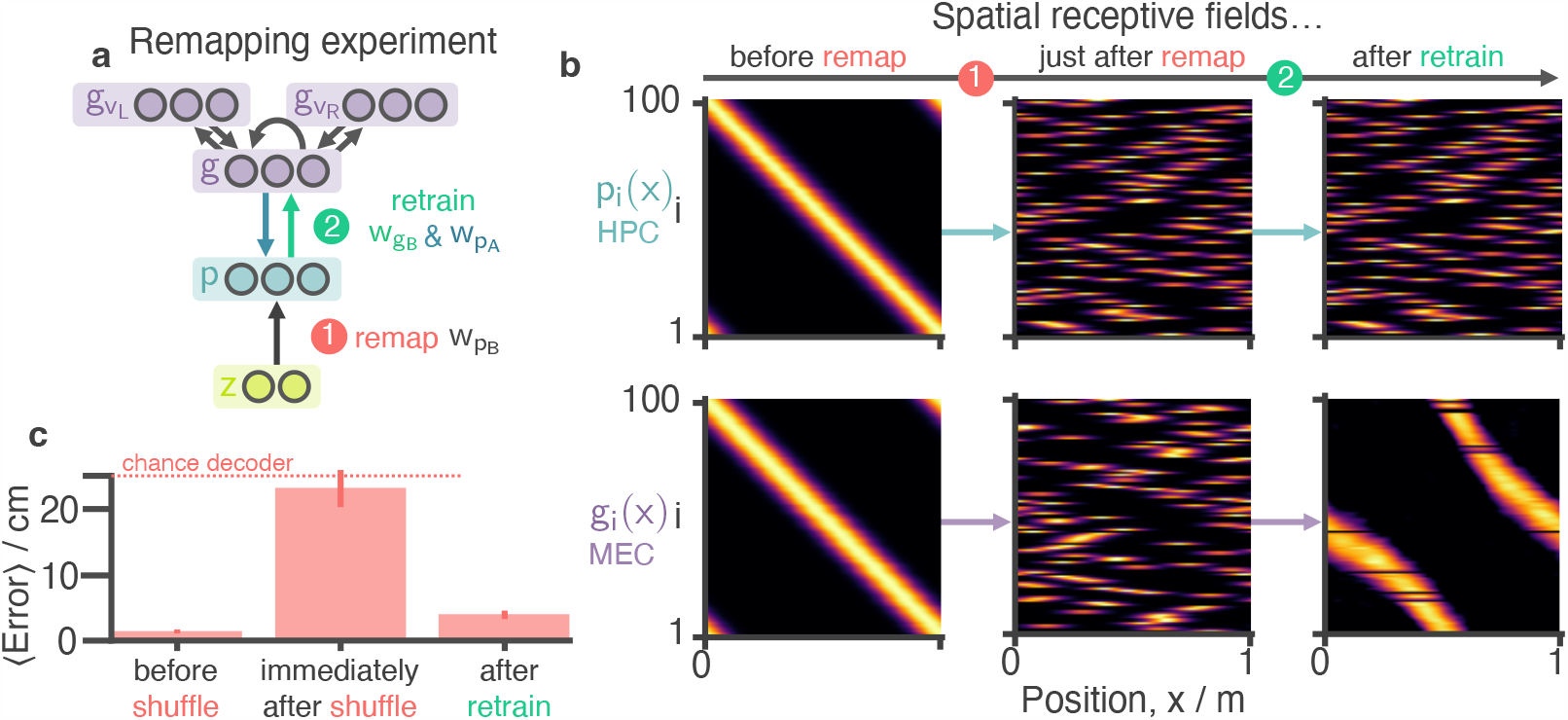
Remapping and transfer of structural knowledge between environments. **a** After training (as in Fig. 2) place cell inputs are shuffled to simulate a “remapping” event observed when an agent moves to a new environment. The agent then retrains for an additional 30 minutes: during this period internal MEC weights, and weights from the conjuctive cells to MEC are held fixed while MEC ↔ HPC weights remain plastic. **b** Recptive fields of the HPC and MEC neuronal populations at different stages in the experiment: Initially after remapping HPC and MEC inputs are randomised. MEC relearns rate maps as they were before remapping but with a constant phase shift. Note: neurons are ordered by the position of their peak activity on the track *before* remapping and this ordering is maintained in subsequent panels. **c** The error (± SEM over 50 trials) after 1 second of path integration is shown at different stages of the experiment. Although path integration is initially disrupted after remapping it recovers despite no relearning of the MEC synapses where the ring attractor is stored.

During biological remapping events place cells remap independently whereas grid cells remap *en masse* with entire modules shifting by the same constant phase [58]. This observation is reproduced in our model: after retraining MEC units regroup with receptive fields as they were before remapping but with a constant phase shift along the track. This re-emergence of structure occurs because the ring attractor seeds a bump of activity on the attractor manifold (during the “sleep” phases of retraining) onto which the shuffled HPC inputs then bind. Since nothing constrains *where* on the circularly symmetric attractor manifold this regrouping can initiate, only relative correlations, modulo a phase shift, are preserved.

Decoding error one second after a sensory lesion is tested just *before* remapping, just *after* remapping and after retraining (Fig. 4c). After the remapping path integration abilities temporarily disappear because the MEC ring attractor is still tuned to the old and invalid HPC receptive fields. After relearning – and despite *no adjustments to the MEC weights*, **w**_*g A*_, *where the ring attractor is stored* – path integration recovers to almost the level before remapping. This differs substantially from other local models of path integration learning [35, 36] which don’t consider plasticity on the ring attractor inputs. In these models, adaptation to a new environment necessarily requires complete relearning of the ring attractor. Instead our model exploits the basic fact that movement (path integration) in one environment is fundamentally the same as in another, one must simply learn a new mapping to/from the ring attractor, “translating” it to fit the new sensory stimuli.

## 4 Discussion

We propose that the hippocampal formation resembles a Helmholtz machine, simultaneously learning an inference and generative model of sensory stimuli. Like previous models [23] medial entorhinal cortex (MEC) sits hierarchically above the hippocampus (HPC) to which it sends generative predictions. Our model differs in the learning rules and neural dynamics: local prediction errors are minimised between distinct dendritic compartments receiving bottom-up and top-down signals. Theta oscillations regulate internal neural dynamics, switching the network between wake and sleep phases. In a navigation task our MEC model forms a ring attractor capable of path integration. Despite simple learning rules and dynamics our model retains key cognitive capabilities of the hippocampal formation including the ability to transfer knowledge across different sensory environments.

Local learning rules are commonly recognised as essential in biologically plausible learning algorithms [43]. However, the importance of learning *scheduling* – how neural systems coordinate or multiplex distinct phases of forward and backward information flow – is often overlooked[59]. Neural oscillations such as theta, hypothesized to temporally coordinate communication between neuronal populations [60], likely play an underexplored role in this regard (neural “bursting” has also been pointed out as a potential solution to multiplexing [61]). One advantage of the wake-sleep algorithm, which this study suggests neural oscillations can support, compared to forward and backward sweeps is that, during convergence, the two phases become highly similar, allowing learning to proceed without affecting perception.

While our discussion has primarily focused on theta oscillations as a mechanism for learning, they have also been proposed as a mechanism for short-range future prediction via so-called “mind-travel”[62]. During the latter phase of each theta cycle (i.e. the sleep phase) gain amplified velocity signals might rapidly drive the MEC activity bump along the manifold allowing the agent to assess nearby upcoming locations. This complimentary proposition could neatly integrate into the framework proposed here and emphasizes the need for further investigation into the multifaceted functions of neural rhythms within the hippocampal/entorhinal system.

Beyond theta oscillations, both faster gamma cycles [63] and the slower physiological states of sleep and wake [64] have been associated with learning. Based on our model we suggest a tentative hypothesis that theta oscillations may be favored due to an optimality criterion; whilst faster oscillations could be a mechanism to prevent extreme drift during sleep that might disrupt learning their frequency might by upper bounded biophysically by the neural time constants associated with the biophysical processes supporting dendritic gating the soma. These ideas, their relevance to other brain regions involved in generative learning, 2D spatial dynamics, and offline memory consolidation/replay remain exciting questions for future theoretical and experimental investigation.

## 5 Supplementary Material

### 5.1 Basic model description

Here we give a general decription of the model. Specifics for each experiment (i.e. learning rates, layer sizes, time constants etc.) are given in later sections.

#### 5.1.1 Dendritic updates

Complete versions of the dendritic update rules (summarised in Eqns (2) & (3)) are given below. We assume dendrites recieve and integrate synaptic inputs according to the following dynamics:

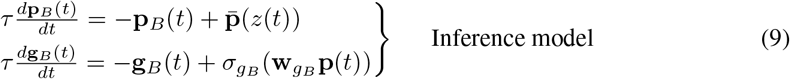

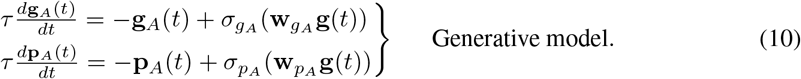

We discretise these dynamics in order to implement them computationally by making the common assumption that neural dynamics are fast (*τ* ≈ 0 ms) relative to the timescale of the synaptic inputs and so the compartments are always at equilibrium, recovering Eqns (2) & (3). This is valid in our regime where the environmental latent updates slowly compared to neural timescales. The notation we’re using admits the possible presence of biases as well as the weights (though biases typically aren’t used) by assuming a row of constant 1’s could be added to the synaptic inputs effectively absorbing a bias into the weight matrix without loss of generality, for example **w**_*g B*_ **p**(*t*) ← **w**_*g B*_ **p**(*t*) + *b*_*g B*_.

#### 5.1.2 Somatic updates

Somatic updates rules (Eqns (4) & (5)) and are repeated here for completeness:

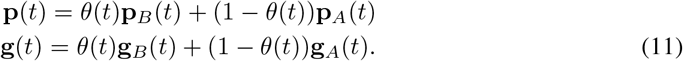

where *θ*(*t*) is a 5 Hz global theta oscillation variable defined by the square wave function:

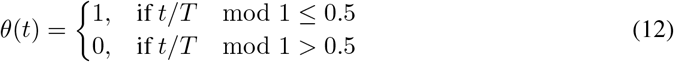

#### 5.1.3 Update ordering

For this hierarchical network of multicompartmental neurons we must specify the order in which we perform these discrete updates to the different layers and the different compartments within these layers. Strictly speaking, when the discretisation timestep *dt* is small this ordering is arbitrary, but we include it here for completeness.

We update the layers from bottom to top: first we update the latent or “environment” and increment the global clock (*z*(*t* + *dt*) ← *z*(*t*) & *t* + *dt*←*t*). Next we update both dendritic compartments of the sensory layer (**p**_*B*_(*t*+*dt*) ← **p**_*B*_(*t*) & **p**_*A*_(*t*+*dt*) ← **p**_*A*_(*t*) noting that it makes no difference in which order these updates are done as they are independent. Then we update the somatic compartment of the sensory layer (**p**(*t* + *dt*) ← **p**(*t*)). Next we work upwards to the hidden layer (**g**_*B*_(*t* + *dt*) ←**g**_*B*_(*t*) & **g**_*A*_(*t* + *dt*) ← **g**_*A*_(*t*) followed by **g**(*t* + *dt*) ← **g**(*t*)) then, if present, the top-most “conjunctive cells” are updated. This gives the following dendritic update rules which are only slightly – and in the limit *dt* → 0, irrelevantly – different from the simplified update rules given in the main text:

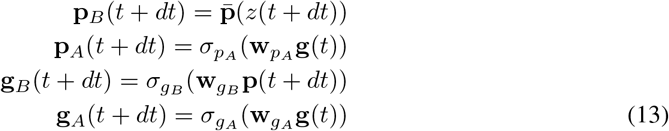

#### 5.1.4 Learning rules

Learning rules are conceptually summarised by the equations given in the main text, Eqn (6). Here we give the *full* equations which include some adjustments to account for the presence of non-linear activation functions and temporal smoothing of the local prediction error learning signals. In our multilayer network all sets of learnable weights follow an equivalent learning rule. For this reason we choose give it here in its most general form: Consider the synaptic weight *w*_*ij*_ connecting from the soma of presynaptic neuron *j* with activation 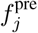 to one of the dendritic compartments of a postsynaptic neuron *i* with activation 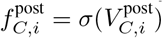 (this could be the basal or apical compartment, *C* ∈ {*A, B*}). Weights are updated on each timestep by an amount:

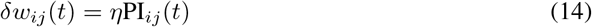

where PI_*ij*_ is (following terminology used in Urbanczik and Senn [43]) the “plasticity induction” variable which is a low-pass filtered measure of the coincidence between the local prediction error and the synaptic input. The prediction error measures how far the activation of the dendritic compartment,

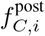, is from the somatic activation 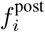. In total, PI_*ij*_ is defined by the following dynamics:

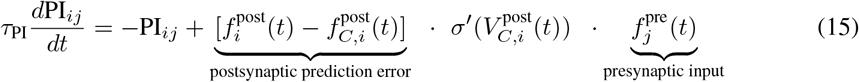

If the prediction error and one of the presynaptic inputs are both consistently large (i.e. over a time period (*τ*_PI_)) then the plasticity induction variable will therefore also be large and the weight connecting the pre- and postsynaptic neurons will be strengthed (thus decreasing future prediction errors). *τ*_PI_ is taken to be the same as used in Urbanczik and Senn [43], 100 ms. Note for fast filtering (*τ*_PI_ → 0 ms) and linear activation functions this reduces to the simplified formulae given in the main text, Eqn. (6).

#### 5.1.5 Synaptic noise

We add synaptic noise to the dendritic activations. Each dendritic compartment maintains its own independent noise variable, *n*(*t*), which is modelled as an Ornstein-Uhlenbeck process. The benefit of modelling neural noise with an Ornstein-Uhlenbeck process is that it is timestep size independent. The dynamics of the noise variable are given by:

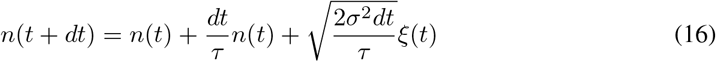

where *ξ*(*t*) ∼ N (0, 1) is a white noise process. These dynamics lead to a stationary distribution of *n*(*t*) which is Gaussian with zero mean and variance *σ*^2^. The decoherence timescale of the noise is *τ* . We fix *τ* = 300 ms and *σ* = 0.01 Hz in order that noise is relatively slow and weak. Noise is added at each timestep to the activation of the dendrites, e.g. **p**_*B*_(*t*) → **p**_*B*_(*t*) + **n**_*B*_(*t*) where **n**_*B*_(*t*).

#### 5.1.6 Measuring the prediction error

Figs. 2b & 3d show the prediction errors of the network layers decreasing throughout training. Here we define how these errors. A consequence of our learning rule is that during wake, the apical dendrites adjust to try minimise the discrepancy between the apical activation and the soma (which, during wake, is equal to by the basal activation). During the sleep phase a short time later the basal dendrites adjust to try minimise the discrepancy between the basal activation and the soma (which, during sleep, is equal to apical activation). If learning is successful we would expect the apical and basal activations to converge, thus we use the following measures of the prediction error to track training performance in both layers of the network:

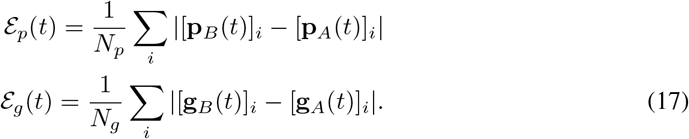

These are then smoothed with a decaying exponential kernel of timescale 60 seconds to remove some of the nosie and better display the learning signal.

### 5.2 Relationship to online Bayesian Inference

Bredenberg et al. [41] derived local synaptic learning rules for a similar hierarchical network performing online latent inference starting from a loss function closely related to the evidence lower bound (ELBO) of variational inference. Here we will not repeat their derivation, instead we intend to highlight their starting point, the most important assumptions they made and the learning rules they derived, finally pointing out how ours differ. The point is to demonstrate that the learning rules we propose are not arbitrary but can actually be derived from a more principled approach to online inference.

Bredenberg et al. [41] consider a network recieving input from a latent variable *z*. The network has two layers, **p**_*t*_ and **g**_*t*_. ^3^ The network is trained to perform online inference over a sequence of observations from the environment, *z*_0:*T*_ . To o this they start rom the loss function

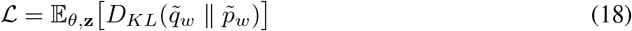

where 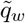 and 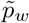 are the following probability distributions over the layer variables **p**_*t*_ and **g**_*t*_:

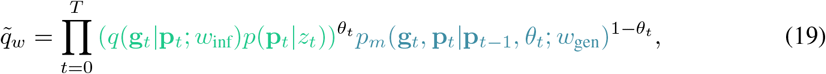

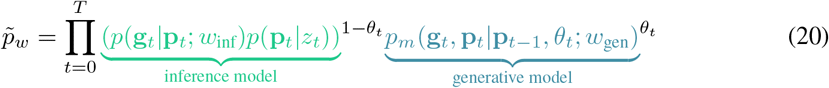

and *θ*_*t*_ ∈ {0, 1} is a binary variable (in their analysis they fix this to oscillate in fixed symmetric phases, e.g. 000111000111…). The two probability distributions, 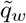 *&* 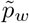, which this loss function attempts to make similar to one another, can be interpreted as the probabilites over the layer variables **p**_*t*_ and **g**_*t*_ in two noisy neural networks^4^ connected as we drew in Fig. 1a: the first network alternates between phases of inference, where information flows bottom up from the latents *z* to the hidden layer **g**, and generation, the opposite (inference-generation-inference-generation…), the second network alternates in exact counterphase (generation-inference-generation-inference…). This loss is a generalisation of the widely used evidence lower bound (ELBO) which corresponds to the case where *θ*_*t*_ = 1 for all *t*. ELBO loss functions seek to make the inference and generative distributions over sensory and hidden variables similar. We will not delve further into the justifications for these types of loss functions other than to state that they are widely used[1].

One of the key conceptual steps taken by Bredenberg et al. [41] (and now us) is to note that processes of performing inference and generation can locally occur simultaneously as long as they are recieved into distinct dendritic compartments. Which dendrite then gates into the soma (i.e. Eqn 4) then dictates the global state (wake or sleep) of the network. It also means, as they show, that the loss can be approximately optimized using local learning rules by comparing the dendritic compartment activation to that of the soma. The learning rules they derive, again translated into our notation, are as follows (note for simplicity we assume all activations are linear since non-linearities add only one additional multiplicative term into their update equations, see equations (14), (15) and (16) in [41]):

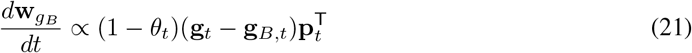

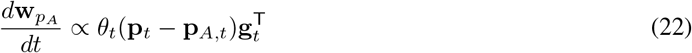

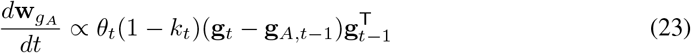

where *k*_*t*_ = (1 − *δ*(*θ*_*t*_ − *θ*_*t*−1_))*θ*_*t*_ is a term which is 1 if and only if *θ*_*t*_ = 1 and *θ*_*t*−1_ = 0 therefore it briefly turns off learning upon switching from sleep to wake.

Reader may like to compare these learning rules to our own as given in the main text Eqns (6). Our learning rules differ from theirs in the following way:

- We relax their discrete time assumption, opting for a continuous time formulation (**p**_*t*_→**p**(*t*) etc.).
- We note that the terms in the equations proportional to *θ*_*t*_ or 1−*θ*_*t*_ which actively turn on or off learning depending on whether *θ*_*t*_ = 0 or 1 are unnecessary since the prediction error term natural falls to zero anyway. For example, in Eqn. (22) when *θ*_*t*_ = 0 the network is in sleep and so **p**_*t*_ = **p**_*A,t*_. In this case the prediction error is zero by definition and learning ceases even without the preceeding *θ*_*t*_ term.
- We disregard the 1− *k*_*t*_ term. Empirically this does not seem to damage our model and theoretically its impact should only be small in our continuous time formulation where the network is only switching from sleep to wake for a negligible proportion of the time.
- Upon provisional theoretical and experimental justification we liken *θ* the theta component of the hippocampal local field potential and set it to 5 Hz.
- Ultimately these changes are surface level. Our learning rules can – and should – be understood as a close approximation to those derived by Bredenberg et al. [41]. Consequently it is appropriate to consider our hippocampal model as learning to perform approximately optimal online Bayesian inference.

### 5.3 Experiment 1: An artifical latent learning task

*N*_*z*_ = 5 independent, autocorrelated, random latent variables are sampled from a Gaussian process with a squared exponential covariance function of width 1 second, samples of these are shown in Fig. 2a and Fig. S2. The sensory layer is large (*N*_*p*_ = 50) relative to the compressed hidden layer (*N*_*g*_ = *N*_*z*_ = 5) and recieves a random mixture of the latents into the basal compartments as described in the text. All activation functions are linear, no layers have biases, all learning rates are set to *η* = 0.01, and the discretisation timestep was *dt* = 25 ms. Weights are initialised randomly [**w**_**g** *B*_ ]_*ij*_ ∼𝒩 (0, 1*/ N*_*p*_), [**w**_**p** *A*_ ]_*ij*_ ∼𝒩 (0, 1*/ N*_*g*_), [**w**_**g** *A*_ ]_*ij*_ ∼𝒩 (0, 0.1*/ N*_*g*_) where the smaller initialisation on the recurrent weights, **w**_*g A*_, was chosen to prevent unstable dynamics.

Before learning – since weights are initialised randomly – basal and apical voltages in the sensory layer are unmatched when tested for a period in wake mode (Fig. S1a). When tested for a period in sleep mode, the small initialisation of the recurrent weights means the hidden layer cannot sustain activity (Fig. S1b, top) which decays and decorrelates rapidly in contrast to the true latents (Fig. S1c). Compare this to after learning where, during wake, basal and apical voltages in the sensory layer are closely matched implying accurate autoencoding through the compressed hidden layer. During sleep, the hidden layer generates sustained activity statistically similar to the true latents (they do not match because during sleep the true latents are not driving the network, even during wake we would only expect our network to represent the true latents in its latent space up to a linear rotation), i.e. its is functioning as a generative model. Note the only source of randomness driving stochasticity and activity in the network is the noise in the dendritic updates themselves.

### 5.4 Experiment 2: Learnable path integration with a hidden ring attractor

An agent randomly moves around a 1 m 1D circular track. The trajectory, *x*(*t*), is sampled using the RatInABox[55] simulation package. This means that velocity is model as an Ornstein-Uhlenbeck process (see Eqn. (16)) with a decoherence timescale of *τ* = 0.7 seconds and a standard deviation of *σ* = 0.5 ms^*−*1^. There are *N*_*p*_ = *N*_*g*_ = 100 neurons in both layers. The HPC dendritic activation function is linear (*σ*_*p A*_ (*x*) = *x*) whilst both MEC dendritic compartments have rectified tanh activation functions (*σ*_*g B*_ (*x*) = *σ*_*g A*_ (*x*) = max(0, tanh(*x*))). Note the choice of activation function means MEC neurons have firing rate 𝒪 (1 Hz). All learning rates are set to *η* = 0.01, the discretisation timestep was *dt* = 25 ms and only **p**_*A*_ *&* **g**_*B*_ have learnable biases.

We model *N*_*i*_ = *N*_*p*_ = 100 inputs which are tuned to the position of the agent according to the following Gaussian tuning curves (these roughly model place cells):

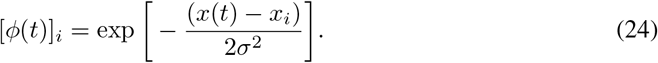

where *x*_*i*_ are centres of the Gaussians evenly spaced along the track. These then linearly drive the basal dendritic compartments of the sensory neurons:

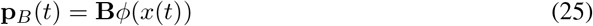

**Figure S1:**
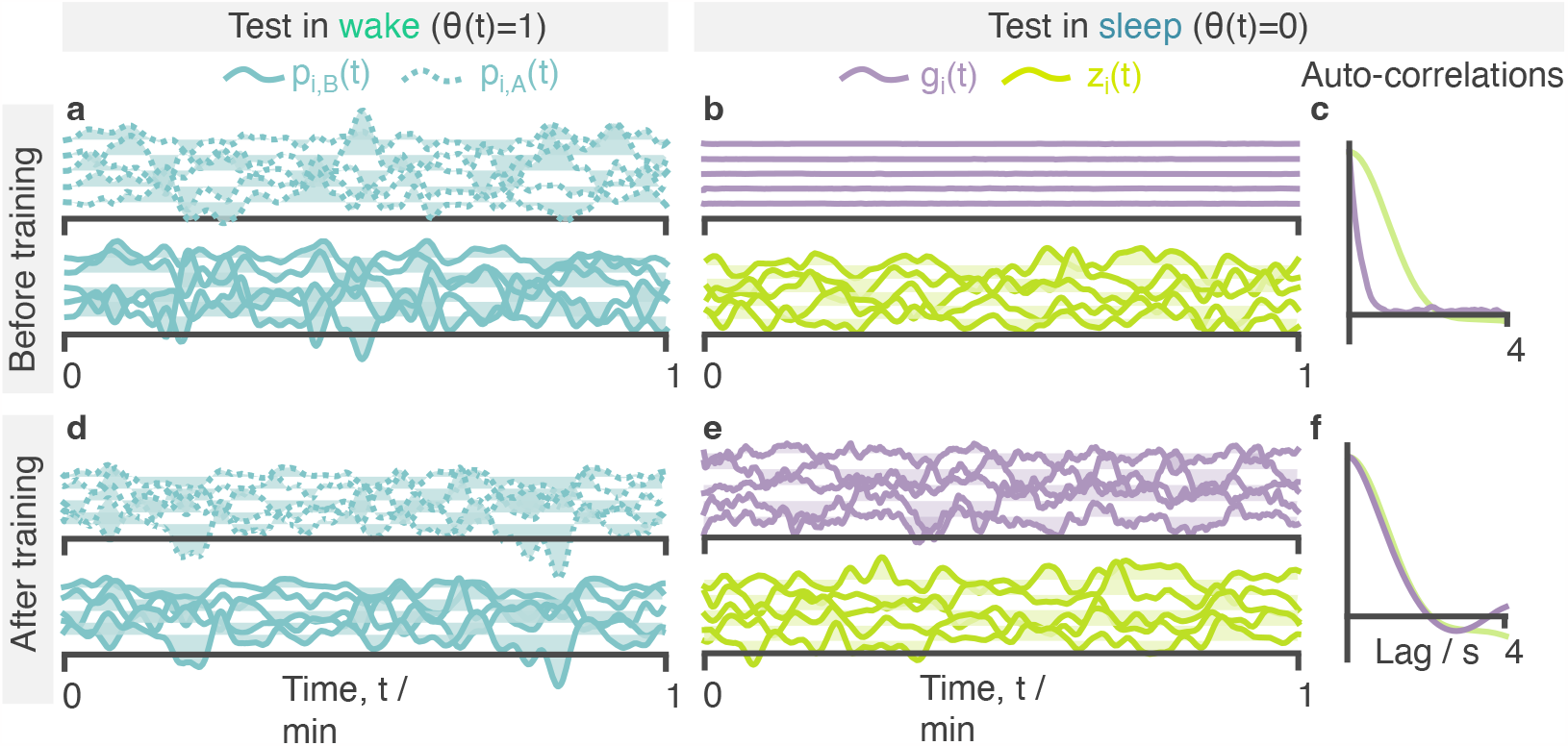
Extended results from the artificial latent learning task. **a** Basal and apical voltages in the sensory layer before learning during a one minute sample in wake mode. **b** Samples of activity in the hidden layer and true latents before training during a one minute sample in sleep mode. **c** Autocorrelations, averaged over the units, for activity in panel b. **d**,**e & f** As in a, b & c but after training.

where, in the results shown in the main paper, **B**_*ij*_ = *δ*_*ij*_ is the identity matrix such that each sensory neuron inherits a unimodel-tuning curve from one and only one of the inputs, i.e. what was stated in Eqn. (8). We show in the supplementary figure S2 that this choice is not particularly critical and the network can learn to perform path integration with random sensory drive 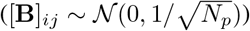.

Velocity inputs are connected as follows: two neurons encode the rectified leftward and rightward velocity of the agent, normalised by the standard deviation *σ*. Note, this means they have firing rates 𝒪 (1 Hz).

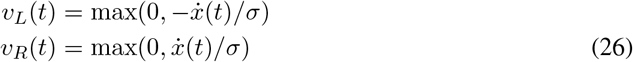

Two sets of conjunctive cells (*N*_*g*_ = 100 in each set) sum inputs from the left and right velocity neurons and the hidden units as follows:

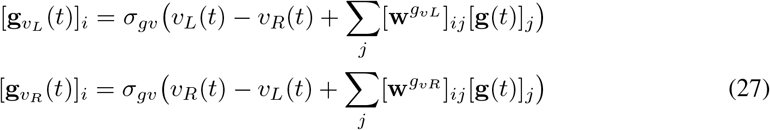

where *σ*_*gv*_(*x*) = max(0, *x*−1) is a ReLU function thresholded at *x* = 1. In the main paper we set [**w**^*gvL*^]_*ij*_ = [**w**^*g*^*vR* ]_*ij*_ = *δ*_*ij*_ so each conjunctive cell is connected to one and only one hidden unit (something we relax in Fig. S2c). The consequence of this connectivity is that a **g**_*v L*_ neuron is above threshold (and therefore active) if and only if the agent is moving to the left*and* the hidden unit it is connected to is active. Rightward motion silences **g**_*v L*_ neurons. Similarly, a **g**_*v R*_ neurons is active if and only if the agent is moving to the right and the hidden unit it is connected to is active. This conjunctive, logic-AND-gate-like tuning to both MEC and velocity is why these neurons are called “conjunctive” cells.

To order the MEC neurons after learning, and thus reveal the ring attractor, we calculate their receptive fields as a function of agent position, **g**(*x*), as though the network is in inference mode (so top-down recurrent connections and drive from the conjunctive cells do not play a role). Then we permute the ordering *i*^*’*^ ← *i* such that the maxima of the receptive feilds move from left to right along the track as the neuron count increases, arg max_*x*_[**g**(*x*)]_*j’*_ *>* arg max_*x*_[**g**(*x*)]_*i’*_; ∀ *i*^*’*^, *j*^*’*^ *> i*^*’*^. The effect of this ordering procedure is shown in Fig. S2, panel a (left hand side, top two panels).

Fig S2a repeats the same path integration test as was shown in the main text Fig. 3 except now we additionally visualise the receptive fields of HPC and MEC (after learning) and show timeseries of both HPC and MEC neurons during the test. Once MEC neurons are reordered by their maxima the ring attractor activity bump can be seen moving up at down the manifold of neurons, even after the sensory lesion. Note again how some MEC neurons have “died” and do not engage in the ring attractor dynamics, forcing the ring attractor manifold to live on the remain subset of MEC neurons.

#### 5.4.1 Position decoding

To quantify the performance of path integration we train a decoder to estimate agent position directly from the HPC population vector. The decoder is trained on positon and activity data from the final 10 minutes of training, after learning had plateaued. The decoder we use is a Gaussian process regressor with a squared exponential kernel, the length scale of which is optimised during fitting. The decoder works well as can be seen in the path integration plots where, before the sensory lesion, the decoded position correctly and accurately tracks the true position.

#### 5.4.2 Robustness of path integration to weight initialisations, plasticity lesions and noise

Since a central claim of our work is that the network can learn, *from random initialisations*, the correct connectivity required to perform path integration, it is important to question where and why weights in our model are not randomly initialised and plastic.

**Sensory weights** The weights from the Gaussian tuned inputs to the HPC sensory neurons, **B** in Eqn. (25), must be non-plastic to prevent the network from rapidly converging on a trivial solution where all input weights fall to zero killing all activity in the network and trivially minimising the local predition errors. They do not, however, need to be the identity function as we chose. Fig. S2 panel b repeats the standard path integration experiment but with a network where 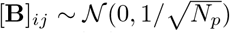, path integration is still learned without any problem. Ultimately this is not particularly surprising since the mapping from the spatially-tuned sensory inputs, *φ*, to the ring attractor in the orginal formulation was already mixed once by the randomly initialised weights from HPC to MEC (**w**_*g B*_ ). This just adds one additional layer of mixing.

**MEC to conjunctive cells** We show in Fig. S2 panel c, that path integration is still learned even when the MEC to conjunctive cell weights are initialised randomly, 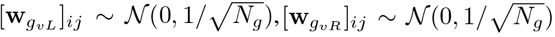 We leave it to future work to investigate this result more thoroughly but comment that it is a notable relaxation on assumptions made in previous models [35, 27] that fine-tuned connectivity from MEC to the conjunctive cells is assumed a priori for path integration (connectivity which would presumably have to be genetically encoded, which seems unlikely). We suspect part of the reason our path integration is robust with respect to the setting of these weights is down to the ability for MEC to construct its own inputs from HPC. This might means the exact form of the activity bump inside the ring attractor can be tailored to fit the specific connectivity to the conjunctive cells – which is perhaps randomly determined during development – in a particular network.

**Plasticity lesions** Path integration, as explored in section 3.2, requires fine tuning the recurrent weights in the hidden layer (*w*_*g A*_ ) and consequently fails when this plasticity is turned off (Fig. S3a). Intriguingly however, we find that path integration does not *strictly* require plasticity between HPC and MEC (as shown in Fig. S3b, echoing results in [35]). However, when such plasticity is removed, the apical input to HPC coming from MEC is unmatched to the sensory input HPC recieves from the environment. As such, any downstream system reading out position from the HPC code would only be able to do so during sleep or wake and not both. This is somewhat restrictive for a system hoping to use the hippocampal formation for online inference and planning. Hence, a primary role of interlayer plasticity between HPC and MEC in our model is to “translate” the environment-agnostic MEC code into the the environment-specific HPC code. This idea is discussed further in section 3.3.

### 5.5 Experiment 3 details: Remapping

To investigate remapping we first train our network to path integrate as described in the main paper. The only difference is that we fix the weights from HPC to MEC to the identity matrix ([**w**_*g B*_ ]_*ij*_ = *δ*_*ij*_ and *η* = 0 on these weights) during this phase of training, this results in MEC neurons with receptive fields equal to those of the HPC neurons (except also passed through a rectified-tanh activation function), Fig 4b left column.

**Figure S2:**
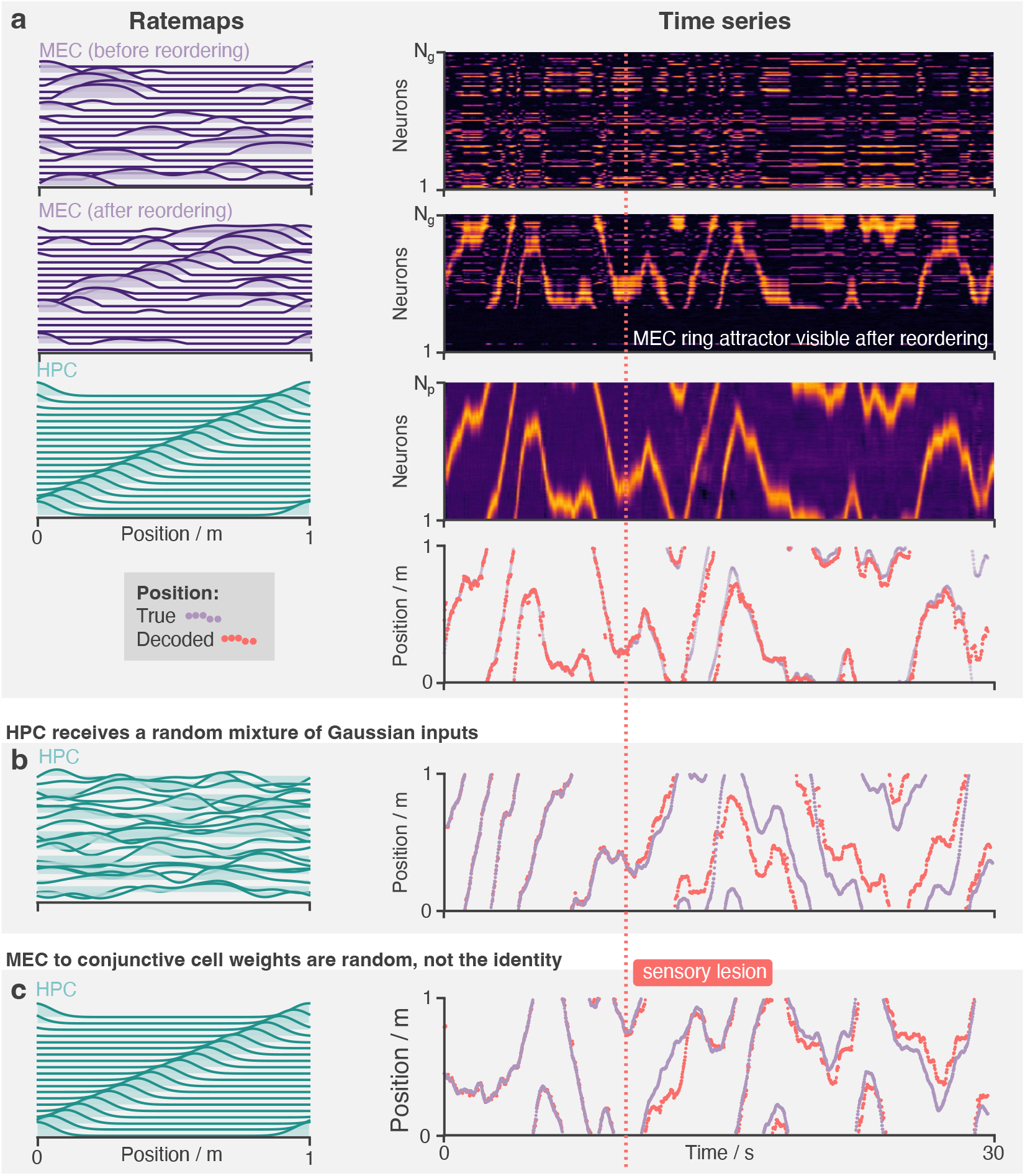
Path integration is performed by a ring attractor in MEC revealed once the neurons are reordered by receptive field peak position. The network learns to path integrate robustly, regardless of the choice of random initialisations. **a** The same path integration test as in the main text is performed here: The top three rows show receptive fields (left) and timeseries activity (right) for the MEC (top two) and HPC layers (third) layers. MEC receptive fields and activity at first appears random. It is only after reordering the neurons by the peak position of their receptive fields that we see the ring attractor manifold. The bottom row shows the decoded position (red) and the true position (purple), demonstrating accurate path integration. **b** Like panel a except, instead of unimodel Gaussian inputs, the HPC neurons recieve a random-sum-of-Gaussian inputs. Nonetheless the network still learns to path integrate (right). **c** Like panel a – with HPC neurons returned to their original Gaussian receptive fields – except in this experiment the hidden units (MEC, **g**) are connected to the conjunctive cells randomly, not one-to-one. The network still learns to path integrate.

**Figure S3:**
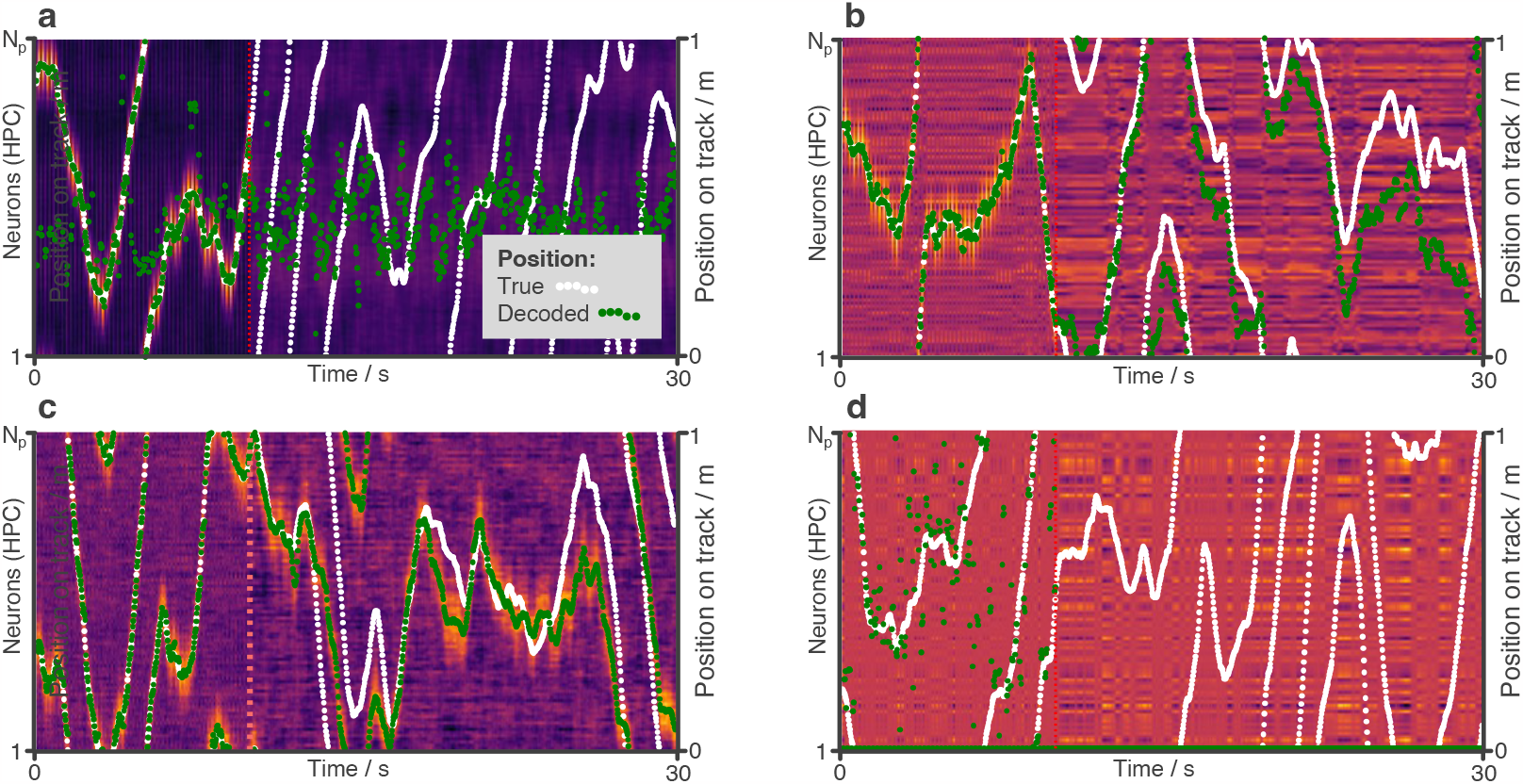
Network response to removal of plasticity and additional noise. The standard path integration experiment is perform and hippocampal activity (as well as true and decoded position) is shown in four modified conditions. **a** Plasticity on the recurrent synapses (*w*_*g A*_ ) is turned off and the network no longer learns to path integrate. **b** Plasticity on all weights between HPC and MEC (*w*_*g*_*B & w*_*p*_*A* ) is turned off. The network still learns to path integrate but inputs to HPC from MEC are not matched to those from the sensory input. **c** Synaptic noise on all synapses is increased by a factor of 10. The bump attractor is now noisier than Fig. 3e but path integration is still accurate. **d** Synaptic noise on all synapses is increased by a factor of 100 at which point learning fails.

In the second phase we begin by randomly permuting the centres of the Gaussian sensory inputs in Eqn. (24). This “sensory shuffle” simulates the sort of hippocampal remapping event which typically occurs when an agent enters into a new environment. The activations of all neuronal layers are reset to zero. A second phase of learning then begins, this time only the weights from HPC to MEC (**w**_*g B*_ ) and from MEC to HPC (**w**_*p A*_ ) are plastic (*η* = 0.01) while the recurrent weights within MEC and the weights from the conjuctive cells to MEC (collectively, **w**_*g A*_ ) are frozen (*η* = 0).

We found that MEC neurons regroup after the shuffle, reestablishing the pairwise correlational structure they had before remapping with, perhaps, a phase shift (Fig. 4b). Once the ring attractor manifold has reappeared in this way the ability to path integrate returns (Fig. 4c). We find these results are clearest when **w**_*g B*_ was fixed to the identity during the initial learning phase as desribed above. Although we don’t investigate this finding thoroughly we suspect it is because the network has an easier time learning the ring attractor since the MEC inputs are already unimodal. With the identity mapping, a tidy activity bump already on the MEC cells before the rest of the ring attractor connectivity is learned, providing a good starting point. Note this matches the standard set up for studies of path integration in, for example, Vafidis et al. [35]). This, perhaps, leads to a ring attractor which is more deeply embedded into the MEC recurrent connectivity structure and which can therefore more easily reestablish itself after a remapping. Nonetheless we discover that MEC is able to relearn a significant portion of the bump attractor structure during the second phase of learning even when this was not the case and **w**_*g B*_ was randomly intialised and plastic during the initial learning, this is shown in Fig. S4. Note how, in contrast to the receptive field shown in Fig. 4b, the MEC neurons are now multimodal and additional bands of correlational structure (in addition to a global phase shift) appear after relearning. We leave it to future work to investigate this further.

**Figure S4:**
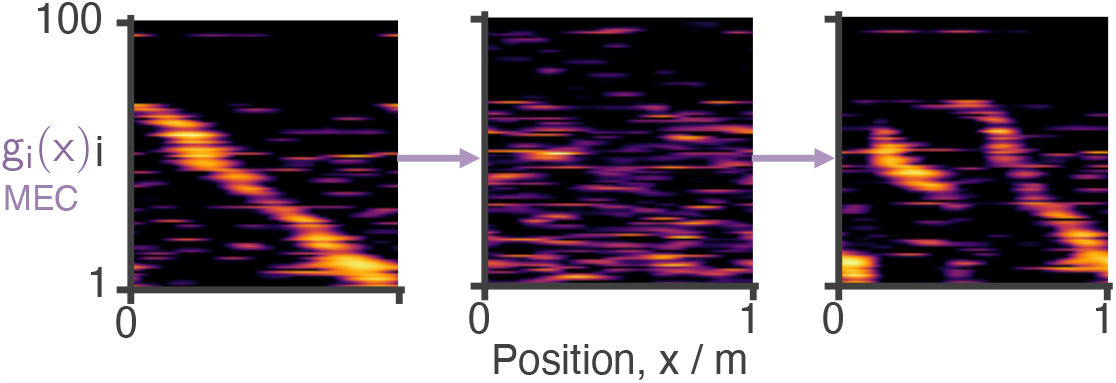
Regrouping of the MEC neurons after sensory remapping but relaxing the constraint that HPC to MEC weights are fixed to the identity matrix during initial learning. This results in MEC neurons with multimodal receptive fields and more complex regrouping dynamics after remapping.

## Footnotes

1 Code provided at https://github.com/TomGeorge1234/HelmholtzHippocampus

2 These labellings conveniently match the notion that inferences are made from layers Below in the sensory hierarchy (bottom-up) whereas generative predictions arrive from Above (top-down).

3 For convenience we have translated their variables into our notation (**g***↔* **r**,**p***↔***s**,*θ↔ λ*, **w***↔ θ*) so it is easier to compare.

4 Note there isn’t *actually* two networks being trained. Instead they use a mathematical trick, deriving from the symmetry in the alternating phase of the theta cycle, to do away with the need to sample from both networks meaning they can deriving local learning rules which can train a single network, e.g. 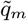, on its own. This single network, like ours, contains both inference and generative models, represented by the two terms in equation (19)

